# Development of intra-oral automated landmark recognition (ALR) for dental and occlusal outcome measurements

**DOI:** 10.1101/2020.12.16.423094

**Authors:** Brénainn Woodsend, Eirini Koufoudaki, Ping Lin, Grant McIntyre, Ahmed El-Angbawi, Azad Aziz, William Shaw, Gunvor Semb, Gowri Vijay Reesu, Peter A. Mossey

## Abstract

Previous studies embracing digital technology and automated methods of scoring dental arch relationships have shown that such technology is valid and accurate. To date, however there is no published literature on artificial intelligence and machine learning to completely automate the process of dental landmark recognition.

This study aimed to develop and evaluate a fully automated system and software tool for the identification of landmarks on human teeth using geometric computing, image segmenting and machine learning technology.

239 digital models were used in the automated landmark recognition (ALR) validation phase, 161 of which were digital models from cleft palate subjects aged 5 years. These were manually annotated to facilitate qualitative validation. Additionally, landmarks were placed on 20 adult digital models manually by three independent observers. The same models were subjected to scoring using the ALR software and the differences (in mm) were calculated. All the teeth from the 239 models were evaluated for correct recognition by the ALR with a breakdown to find which stages of the process caused the errors.

The results revealed that 1526 out of 1915 teeth (79.7%) were correctly identified, and the accuracy validation gave 95% confidence intervals for the geometric mean error of [0.285, 0.317] for the humans and [0.269, 0.325] for ALR – a negligible difference.

It is anticipated that ALR software tool will have applications throughout Dentistry and anthropology and in research will constitute an objective tool for handling large datasets without the need for time intensive employment of experts to place landmarks manually.

## 1. Introduction

Dentistry, like many other medical disciplines, is moving towards a more digital approach, using technology to the advantage of higher quality diagnostic and therapeutic patient care. Digital imaging and 3D imaging reconstruction play a greater role than ever in both diagnosis and treatment planning in a range of disciplines and applications in the fields of medicine and dentistry. The challenge is to utilize the developments in digital imaging and computing technology to create software to automate and improve certain procedures such as diagnostics and outcome measurements.

In the dental field, clefts of the lip and palate (CLP) are orofacial conditions with many different sub-groups and of varying severity. The sub-phenotypes are accompanied by complications from birth which include feeding and swallowing difficulties, speech impairment, hearing problems and dentofacial growth and development. The last of these, facial growth disturbance is an important outcome as it affects the quality of life has a significant psycho-social impact, especially during adolescence (1, 2) and it is important that it is measured accurately and objectively if possible.

Ongoing research in the field of CLP continues to determine the optimal timing in relation to oro-facial development for primary surgery, and which surgical method(s) should be employed. Robust and reliable assessment of treatment is an essential part of modern clinical governance, to allow for the development of better surgical protocols for those treated in the future. One key ‘core’ outcome measure of primary surgery of the palate is facial and maxillary growth and this is measured by the degree of maxillary arch constriction which may be affected by a range of factors including severity and surgical technique. Several scoring systems have been developed to quantify the degree of maxillary arch constriction by examination of the dental arches relationships. Some scoring systems such as the GOSLON (3) and 5-Year-old index (4) are subjective, categorical and require examiner calibration and elements of clinical judgement, while others are designed to be objective and quantitative e.g., the Modified Huddart Bodenham (MHB) index (5) (Figure 1).

**Figure 1:**
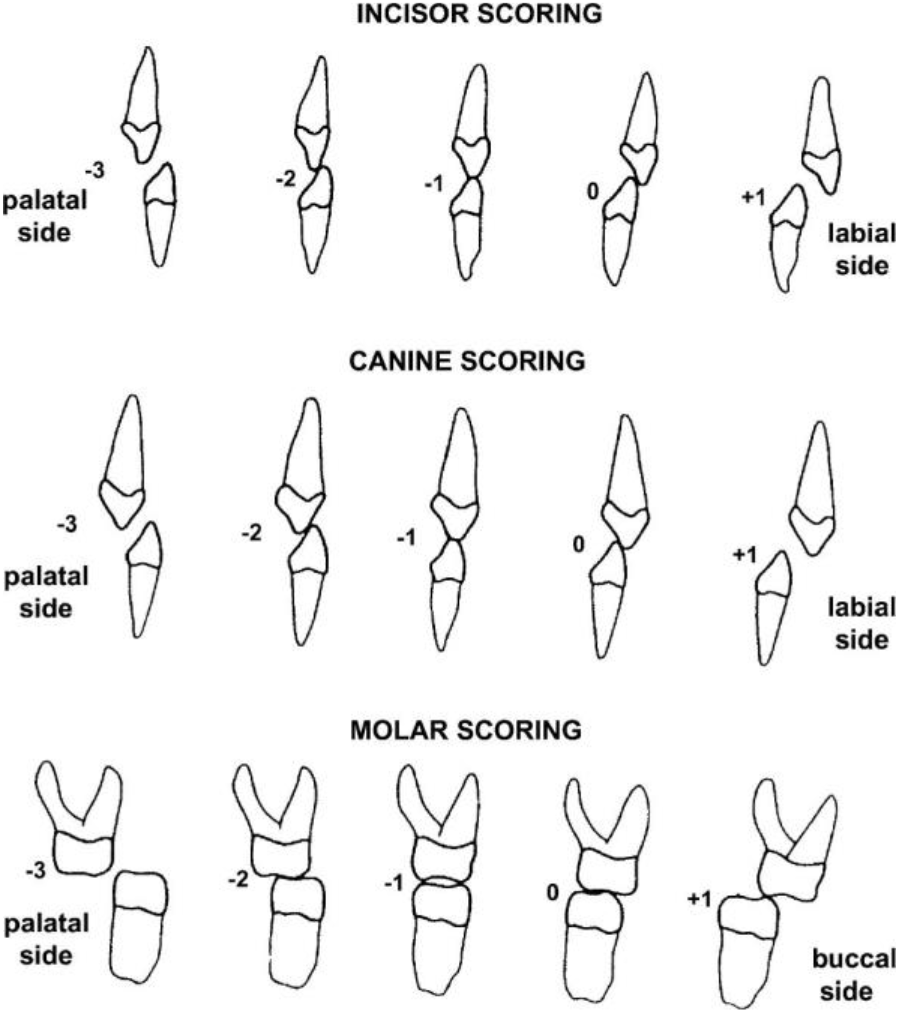
The modified Huddart Bodenham (MHB) scoring system for CLP outcome measurement

The latter system where scoring is quantified using an ordinal scale lends itself to digital analysis, and the scoring has been shown to be suitable for fully automated software (6). Furthermore, it can be used for all cleft types, does not require calibration, is systematic and is objective ((Figure 1). The MHB scoring system has also been validated on plaster models, photographs and digital models that can be captured using intra-oral scanning. (7, 8)

The process of optical scanning to produce digital impressions is the preferred technique when compared to traditional impression taking as this is more comfortable for the patient (9). Intra-oral scanners produce a 3D model of the scanned arch that can be saved and analysed on a computer, and in addition to many other advantages, they do not require physical storage space, making it easier for clinics to store and archive patient data for clinical and research purposes (9).

A useful additional feature of the digital models is that they lend themselves to computer manipulation, and automated data analysis techniques to produce objective outcome scoring tools. In the field of CLP outcome measurement using MHB on digital models, a semiautomated software tool programmed by Rhino 5C++ SDK was developed by (10). This enabled objective scoring, eliminated the need for calibrated examiners and anchor study models and was found to be accurate and reliable. However, it still requires the manual identification of landmarks on the dental arches which is time consuming and is adversely affected by human error.

In developing automatic scoring software, accurate landmark identification is crucial and currently landmarking dental tissues is manual. And so, despite the potential benefits that come with the 3D information capture, this new technology also introduces new challenges, in data capturing, data manipulation and efficiencies in time and labour saving. (11, 12) This is particularly so when the analyses of multiple cases are required, despite its reliability, manual handling becomes impractical and difficult to complete. Manual identification has been prone to errors such as landmark inaccuracies and varying degrees of subjectivity from different professionals thus making it hard to cross-compare and analyse (13). This motivates the importance of automatic landmarking and an automatic scoring software based on it, and places significant value on the development of an automated software tool that is based on automated landmark recognition (ALR).

Software for automatic landmarking and for an objective outcome measure using the MHB scoring system is required. This would replace the need for time-consuming manual scoring and eliminate human error and revolutionise outcome-based cleft research, ultimately resulting in an objective outcome measure of primary surgery. The aim of automated scoring is increased efficiency, standardisation of the scoring of surgical outcomes allowing inter-centre comparisons of treatment protocols and the ultimate aim is to produce objective evidence of where and how improvements in standards of care can be achieved for individual cleft phenotypes.

The overall aim of this study was to create and validate the accuracy of a fully automated ALR algorithm and to use MHB as an exemplar outcome measure, and in the process to test the following null hypothesis. “ALR as used on digital models for MHB scoring is no less accurate than manual landmarking”.

## 2. Materials

Dental models (see Table 1) processed by the ALR software are represented in the STL format, a common open-source 3D file format. The models were acquired with the use of an intra-oral scanner, either directly from the patients mouth or indirectly, by scanning patients’ study models created after taking conventional impressions. Within an STL file, only the surface of the object is described. STL files contain no colour or texture information. While STL files generally contain no scale information and the units are arbitrary, scanned dental arches are measured in millimetres.

**Table 1:** Types and counts of the 239 scanned models used in this study

| Count (n) | Arch | Type | Dentition | Characteristics |
| --- | --- | --- | --- | --- |
| 24 | Maxillary | Digital model | Permanent | Orthodontic subject |
| 21 | Mandibular | Digital model | Permanent | Orthodontic subject |
| 16 | Maxillary | Intra-oral scan | Permanent | Forensic research archive |
| 17 | Mandibular | Intra-oral scan | Permanent | Forensic research archive |
| 81 | Maxillary | Digital model | Primary | 5-year OFC audit archive |
| 80 | Mandibular | Digital model | Primary | 5-year OFC audit archive |

Typically, the surface of the object displayed is referred to as a mesh. A mesh is hollow and is constituted by small triangles. The constituents of the mesh are defined by the (X, Y, Z) values of its three vertices. In the initial feasibility studies and experimentation on automation of landmark recognition on various dental anatomical features in both maxillary and mandibular arches and various characteristics of dental occlusion, a wide range of models were used. Forty five were digital images from scanned plaster study models of permanent dentitions of subjects with a variety of malocclusions who attended the Orthodontic department of Dundee Dental Hospital, 24 were maxillary arches and 21 mandibular; 33 were derived from direct intra-oral digital scans that were recorded for a forensic odontology research project, and were also of permanent dentitions, while 81 sets were from a 5 year old cleft lip and/or palate cohort that were recorded for audit and research purposes. All subjects whose models were used had given their consent for the use of their models for research purposes and all were anonymous at the point of access to the researchers. The upper and lower mismatch can be explained by an occasional model being unusable or unavailable. The groups and number of models per group are listed in Table 1.

### Methods and materials

An algorithm was developed to convert the landmarks identified by ALR in 3D space to a horizontal measurement of the relationship between the maxillary and mandibular dental arches. The algorithm is broken down into steps. The flow diagram in figure 2 gives an overview of the process.

**Figure 2:**
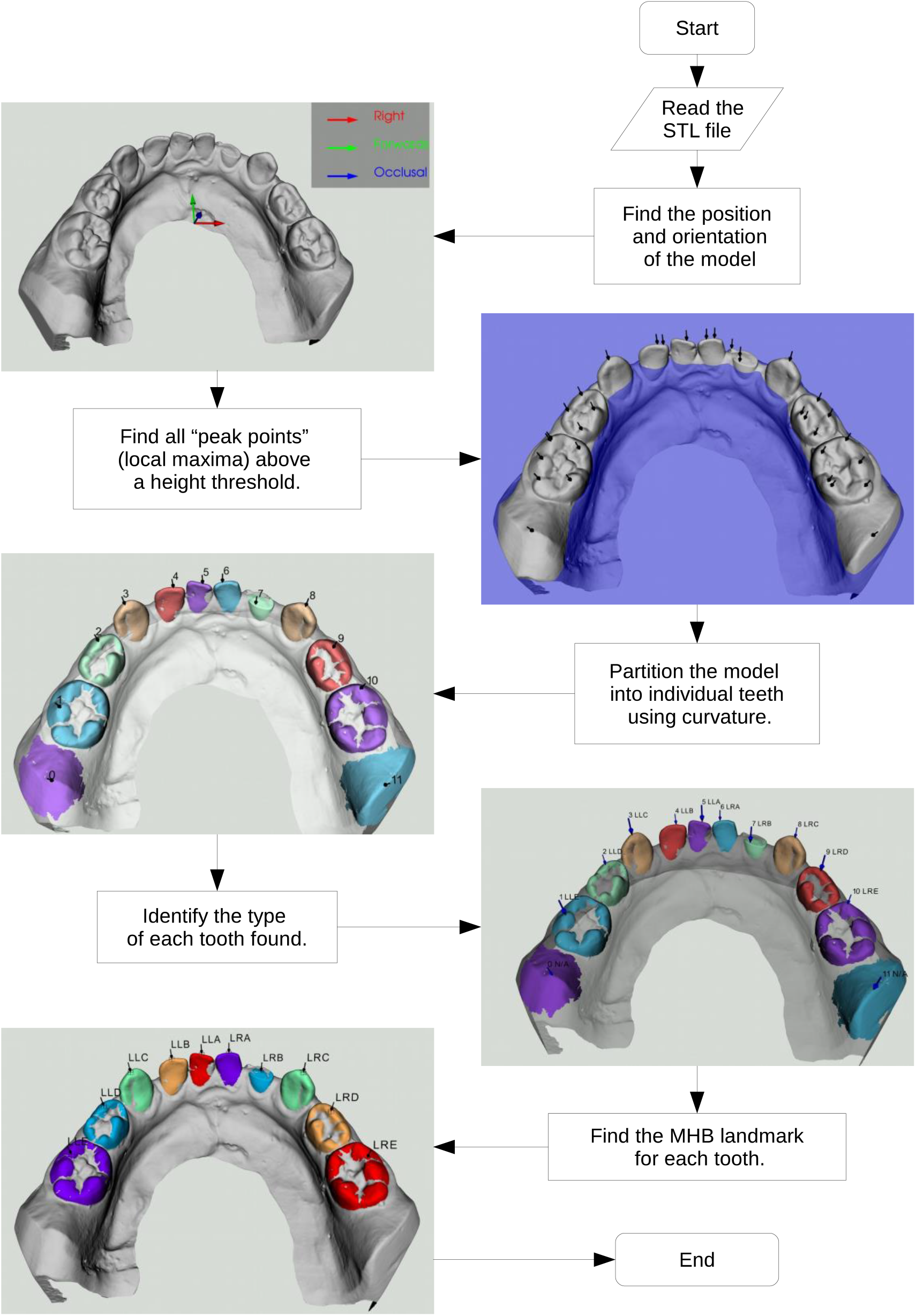
The stepwise process used for automated landmark recognition (ALR) in the deciduous dentition.

Before analysis of the file, the software specified whether the model represented mandibular or maxillary arch and deciduous or permanent dentition. The ALR software then enabled the automatic recognition of the teeth using the information from the STL filename.

### Orientation

Because different scanners use different orientation conventions, the mesh’s orientation needs to be registered by the software to ensure accurate analysis. The appropriate orientation is calculated primarily by using the *Principal Component Analysis* (PCA) method. PCA checks the covariance from the centre of mass in all directions. It provides unit-vectors to represent left, right, backwards, forwards, down, up and occlusal. The vertical directions (up, down, occlusal) need to be particularly accurate, whereas horizontal directions can be more approximate. In simple terms, PCA finds the shortest, middle-length, and longest dimensions of the object.

A digital dental model is wider (transverse direction) than it is long (antero-posterior dimension) and longer than its height. Accordingly, the output of PCA on a dental model should yield transverse as the principal (longest) dimension, anterior or posterior as the second and vertical the third. The vertical directions are further refined by fitting a plane to the tooth tips. PCA’s key advantage is that the model can be positioned and orientated anywhere and PCA will track it as easily as if the model were already positioned ideally.

### Peak points identification

The next step is to identify all the “*peak points*” (local maxima) of the mesh. As mentioned earlier, the landmarks that need to be identified are located on the tips and cusps of teeth, which tend to be the highest parts of teeth. Mathematically these can be translated as local maxima in the occlusal direction. These highest points will be referred to as “peak points” or “peaks” (see Figure 3). In this stage, peak points are identified, but at this stage it is not clear yet what each peak point represents. To reduce the number of non-tooth features found, only the top (occlusal) part of the model is searched. A height threshold of 6mm below the highest point is used to avoid irrelevant information being included. This threshold is low enough to capture teeth that are worn down or only partially erupted. Some points identified will be of no importance, and some may be duplicated, but they are filtered out by the software at a later stage by more sophisticated means. The successful completion of this step simply requires the software to identify at least one peak point per tooth on the mesh.

**Figure 3:**
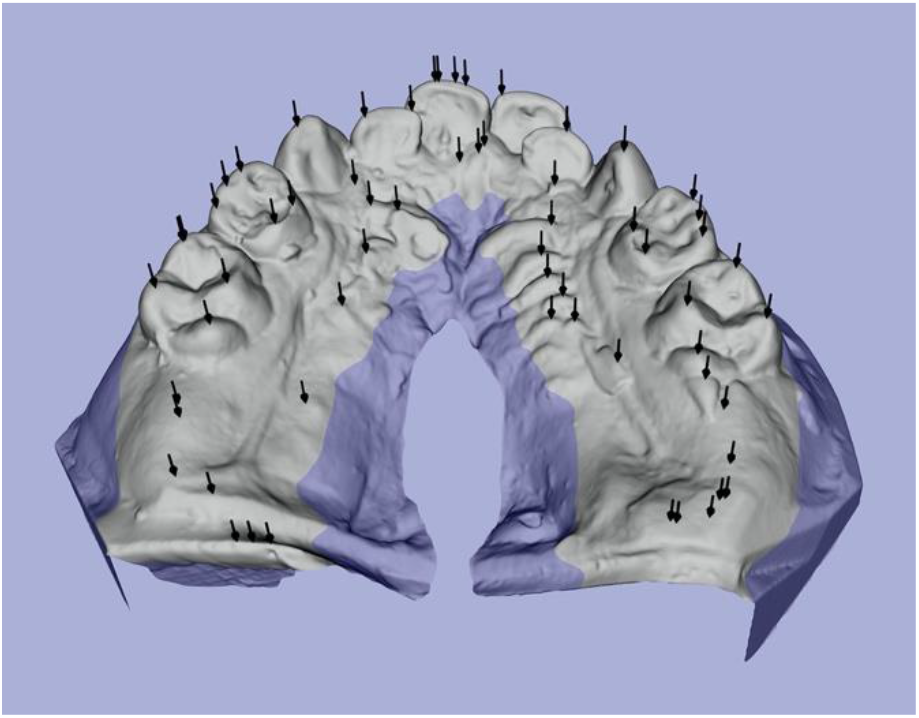
Peak points found (black arrows) and height threshold (blue) on dental models.

### Tooth partition

The software then *partitions* the model into individual teeth. The identification of individual tooth boundaries is based on curvature. The curvature value of a particular surface point is a quantitative measure of its deviation from a flat surface. The definition of curvature in the ALR software uses positive values for outside corners (a bump, cap or tip) and negative values for inside corners (a slot, groove or crease). The joint between each tooth and the gingivae is a crease and therefore the curvature along it is negative. The software starts the scan at the top of a tooth and recursively includes adjacent mesh triangles until an edge of considerable negative curvature is detected, indicating the boundary between tooth and gingivae. The region covered will be referred to as the starting point’s area. Each peak point found in the previous step is used as a starting point for curvature analysis. Peaks included on the same tooth will have areas that overlap. By testing for and merging overlapping areas, one tooth is identified and duplication of teeth in the software is avoided. Any detected peaks that were not located on a tooth will not be bounded by the creases of tooth-gingival joints. As a result, the software might incorrectly include a large part of the model as a single partition. By imposing the rule “stop if travelled more than a tooth’s width away from the starting point”, and testing if that rule was actually used, non-tooth peaks and their corresponding areas can be identified. These non-tooth areas are labelled “spilled” and are discarded.

The scenarios above are demonstrated in figure 4. The simplest case is the LR3 (green) with one peak on one tooth meaning no further work is required. The red dot is a *spilled* peak, it encountered no tooth boundary below it and would have tried to fill most of the base without the maximum traversal rule. Multiple peaks are on the LL5 (cyan) but because their areas overlap, they can be merged thus avoiding duplicity. The buccal (blue) and lingual (yellow) halves of the LR6 don’t overlap but are at the same point round the jawline and can therefore be merged.

**Figure 4:**
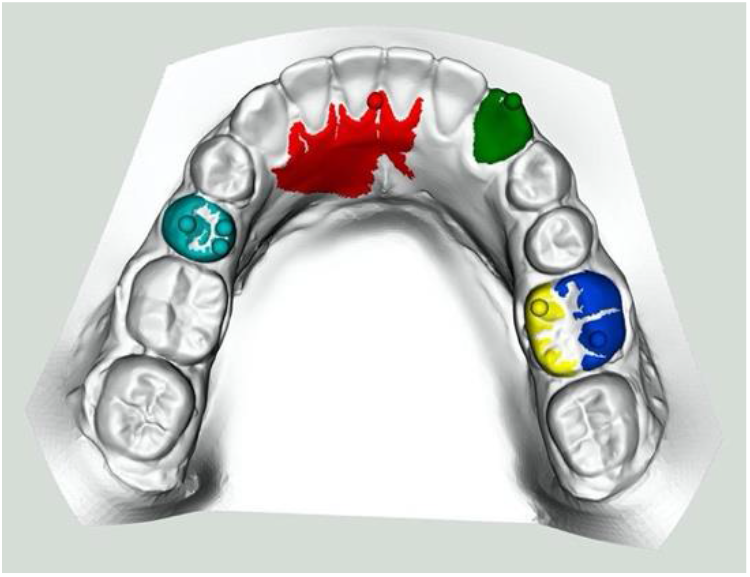
A few different tooth partition scenarios demonstrating peaks (balls) and their areas on a dental model of the permanent dentition.

### Tooth assignment

The successful completion of this step results in the partitioning of each tooth, without any duplication and with almost all the non-tooth structures removed. At this stage, no information regarding the delineation of each tooth is available, and the next step is tooth assignment. The partitions identified within the mesh by the software will be referred to as blobs. Each blob represents either a whole tooth, a mesial or distal half of a molar, or in rare instances a non-tooth feature. The software sorts and enumerates the blobs from left to right round the jaw. The end goal and final output of the tooth assignment method is to correctly label each originally unknown tooth with the appropriate tooth type. In labelling the tooth types, the software uses machine learning techniques on various tooth characteristics, such as total surface area. Each unknown blob in the model is compared to a gold-standard database of manually labelled blobs (training set, see figure 5). The tooth characteristics cover all tooth types and yield single value per blob. The objective is to minimise the total mismatch between the characteristic values of each blob with the training values of the tooth type it is assigned. There is no limit to how many characteristics can be used, and in general it was considered that the more the better. It transpired, however, that only a few basic dimensional measurements are needed to give robust results.

**Figure 5:**
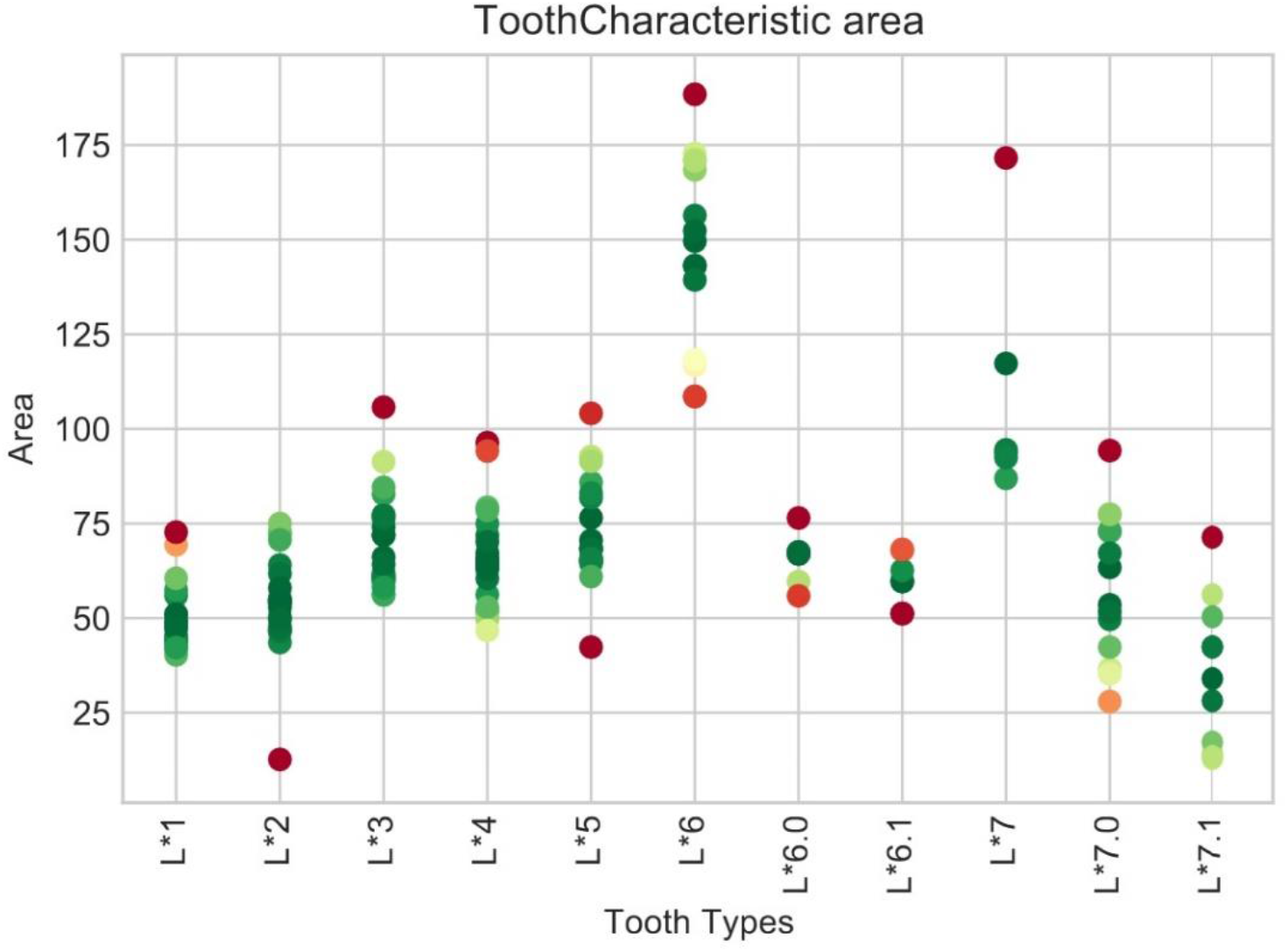
Surface area tooth characteristic chart showing surface areas from the training set by tooth type during tooth assignment.

The tooth types searched for are referenced using Palmer shorthand (e.g., UR2 for an upper right 2nd incisor) with a slight extension to include half-molars. UR6.0 and UR6.1 represent the mesial and distal halves of an UR6, respectively. The software generates a list of potential tooth types based on the model’s jaw type (mandibular or maxillary) as specified at the beginning of the process.

After tooth assignment, the ALR system is then primed for use on whatever system it is applied to. In this context, it was applied to the measurement of maxillary arch constriction in a cohort of subjects who had surgical correction for cleft lip and palate. To do this the points on maxillary teeth and mandibular teeth were mapped and the buccal segment teeth crossbites and incisor overjets were given Modified Huddart Bodenham (MHB) scores.

### Validation

Finally, the software validation was conducted by comparing ALR landmark placement, to landmark identification of the same models manually by dental professionals, using comparison to the consensus x, y, z coordinate of each landmark in 3D space.

During the validation process, 20 models (10 pairs of mandibular and maxillary arches) were scored by the ALR system. The same models were also scored by professionals in the field of dentistry individually after they were given the same instructions. The results of the validation process are displayed in the results section (Table 2 and Table 3) and analysed in the discussion.

**Table 2:** Qualitative evaluation results during the process of tooth partition and assignment as raw per-tooth counts and percentage ratios for adult and deciduous teeth.

|  | Cast Adult |  | Intra-oral Adult |  | Cast Deciduous |  | Overall |  |
| --- | --- | --- | --- | --- | --- | --- | --- | --- |
| **OK** | 475 | 80.8% | 157 | 74.8% | 894 | 80.0% | 1526 | 79.7% |
| **Partition Error** | 46 | 7.8% | 28 | 13.3% | 136 | 12.2% | 210 | 11.0% |
| **Wrong Assignment** | 67 | 11.4% | 25 | 11.9% | 87 | 7.8% | 179 | 9.3% |
| *Wrong Kind* | 22 | 3.7% | 9 | 4.3% | 32 | 2.9% | 63 | 3.3% |
| **Total** | 588 |  | 210 |  | 1117 |  | 1915 |  |

**Table 3:** Average distance in millimetres from the *consensus* coordinates from each of the 3 examiners (A, B and C).

| Deviation (mm) | Software | Humans |  |  |  |
| --- | --- | --- | --- | --- | --- |
|  |  | A | B | C | Mean |
| **Incisors** | 0.454 | 0.565 | 0.387 | 0.349 | 0.434 |
| **Canines** | 0.369 | 0.630 | 0.322 | 0.291 | 0.414 |
| **Premolars** | 0.402 | 0.472 | 0.313 | 0.296 | 0.360 |
| **Molars** | 0.334 | 0.382 | 0.269 | 0.310 | 0.320 |
| **Overall** | 0.389 | 0.517 | 0.310 | 0.300 | 0.376 |

## Results

After the creation of the software, it is important to be able to quantify its performance so we can assess the level or reliability. During the refinement process for the ALR, after arch orientation, tooth partition and tooth assignment were conducted, and the quality of each ALR step was assessed separately to ensure validity of the final landmarking.

A qualitative assessment of the software’s ability to identify teeth was performed on all the models listed in table 1. Teeth were labelled by hand and by the machine and compared per tooth. Table 2 provides summary statistics as raw counts and as percentages. To summarise the row headers: *OK* means the software found and correctly labelled the tooth. *Partition Error* means that the tooth was not found (see subsection Tooth partition) Wrong *Assignment* means that the tooth was found but mislabelled (most of which were lower incisors) (see subsection Tooth Assignment). *Wrong kind* is a subset of *Wrong Assignment* where the software also chose incorrectly from incisor, canine, premolar and molar. In addition to the mistakes above, 1 deciduous model failed at the *orientation* step. The *find peaks points* process has provided accurate results every time.

For the ALR validation, 20 cast models (213 teeth) were processed automatically by the software and manually by three experts to independently identify the assigned landmarks. The results of the landmark accuracy validation, comparing the coordinates of landmarks placed by experts to landmarks placed by the machine, are shown in Table 3. When comparing the coordinates of the landmarks indicated by the software and the mean coordinates of the manual examinations, the results are similar, with similar levels of deviation from consensus landmarks.

As demonstrated in table 3, the overall landmark placing of the software averages out at 0.389mm and the landmark placing by humans at 0.376mm. This represents very accurate landmark placement by the software with the mean difference being 0.013mm. That table also shows that the results have more variation between manual landmark placement by different users, which is currently the gold standard. So, the software placement of the landmarks will have the same diagnostic value and will eliminate the intra-rater variations.

The 95% confidence intervals for the geometric mean error of the humans are [0.285, 0.317] and for ALR. are [0.269, 0.325]. The difference between humans and ALR shown in Figure 6 is negligible.

**Figure 6:**
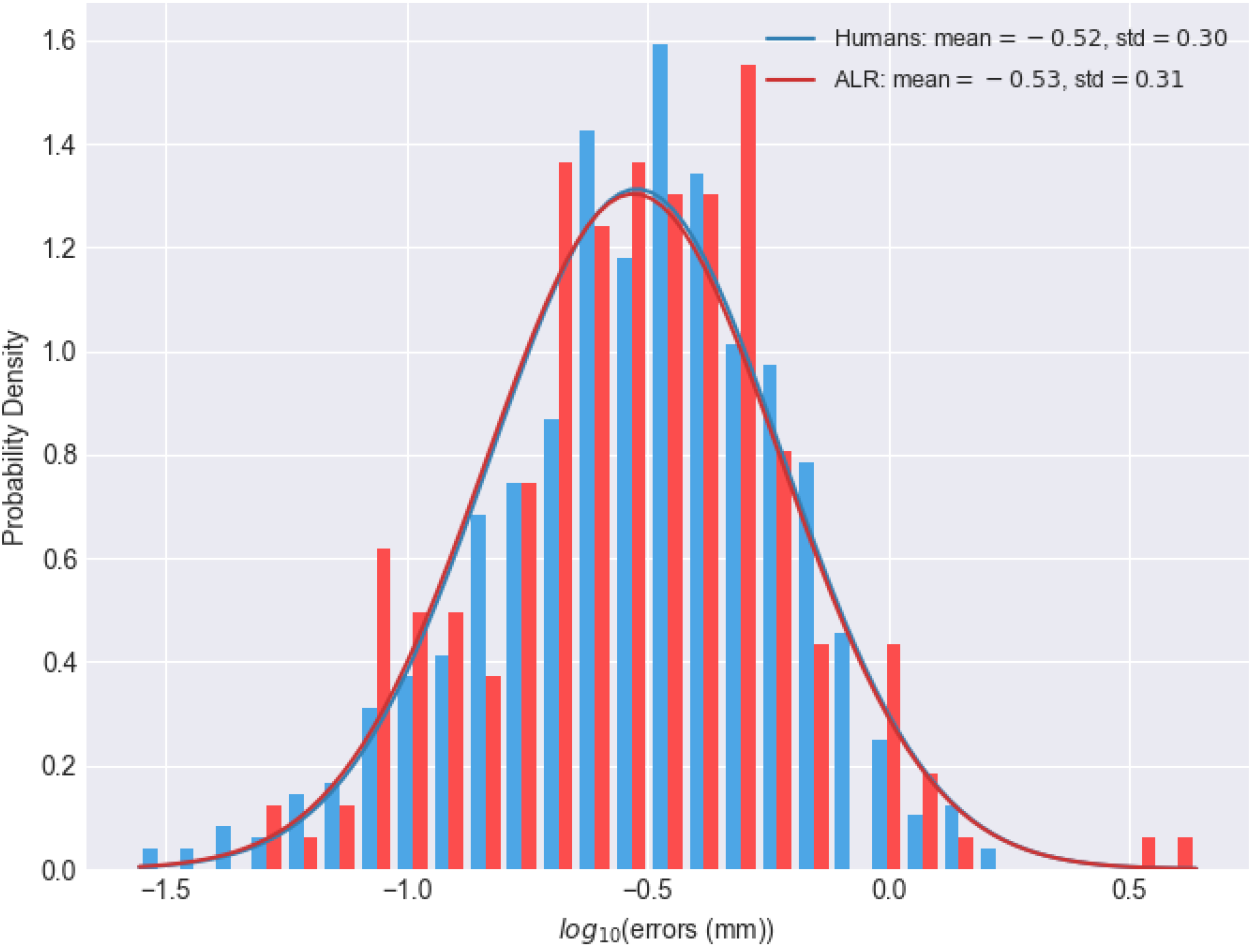
The distribution of placement errors (bar graphs) from the humans’ and ALR’s landmarks and best fit Gaussian distributions (lines) for each.

## Discussion

The work described in this paper builds on the successful development of a plug-in on an open 3D development platform digital technology by Ma et al, in 2017 (10) using Rhinoceros software, version 5. Clinicians selected pre-defined landmarks on cusps of mandibular and maxillary teeth on 3D digital models. A three-dimensional cubic spline was used to generate a mandibular curve, and a best-fit horizontal mandibular reference plane was produced using a least-squares method. Their project was aimed towards an automated scoring of Cleft Lip and Palate outcomes and used horizontal distances projected from the shortest three-dimensional distances between the maxillary cusps and the mandibular curve to calculate the modified Huddart and Bodenham (MHB) score. This automated scoring of digital models using the MHB system produced similar results to manual scoring by experienced experts, the current “gold standard” thus validating its use.

However, one major drawback remained, the fact that the visual and manual identification of the dental landmarks that are selected to enable the MHB score to be allocated still required operator intervention. The Ma et al., development (10) was proof of concept that the digital system with an MHB algorithm was a valid, reproducible and accurate substitute for manual scoring. The ALR software described here aims to overcome this drawback by removing the element of human error through machine learning and thereby improve both diagnosis and assessment of treatment outcomes across many fields of dentistry. It facilitates research and data collection, improving speed, accuracy and standardisation of data analysis, beyond what is feasible with manual methods, and is particularly suited to “big” data.

One of the main challenges faced during the development of ALR was training the software to discriminate between different tooth types. There were several particularly hard to distinguish types, some of which we have overcome, giving a good degree of discrimination, but others such as discrimination between first and second premolars still need improving. The machine learning model is designed to be easily extendable and could later include other known (to humans) distinctive properties which would greatly improve this. A larger training dataset would likely help as the model should be usable as an outcome measure with smaller as well as larger datasets.

Another drawback is that ALR only supports exclusively deciduous or permanent dentition. One of the next additions to the software would be to enable support for mixed dentition. The barrier to this feature lies solely in the Tooth Assignment section. One area that still requires work is making the ALR software more user-friendly. Currently, only parts of ALR even have a user interface. A simple, intuitive user interface, not requiring extensive training, will lead to a reduction of human error in using the software.

ALR is an example of a technological advancement made possible by collaboration of Dentistry and Mathematics, it is a valuable tool that can change the way that many dental related problems are approached. The automated system facilitates quicker and more reliable outcome assessments by minimizing human errors. By standardizing outcome assessment in cleft care, multi-centre comparisons for audit and research can be simplified, allowing centres throughout the world to upload three-dimensional digital models or intraoral scans of the dental arches for remote scoring. Thereafter, these data can feed back into the global database on orofacial clefting as part of the World Health Organization’s international collaborative “Global Burden of Disease” research project for craniofacial anomalies.

Validation of the ALR software means that this can be used to identify landmarks with greater consistency than human identification. ALR enables analysis of large quantities of patient models to provide more accurate predictions for different therapies. That will result in the creation of more specific guidelines that will facilitate the best possible results for the patients, so they can have the best quality of life they possibly could. The fact that ALR obtained results will always be consistent will make collaboration and cross comparison between researching bodies across the world possible without expert clinicians having to travel from site to site, making such research faster, cheaper and easier. The method and software are found to be a valid, reproducible and consistent for identifying landmarking points and through algorithms providing for the first time a fully automated system.

## 6. Conclusion

This paper is the first report to describe the development of an automated dental landmarking and scoring software via a combination of geometric computing, data fitting, tooth image segmenting and machine learning. This fully automatic landmark recognition can be adapted for a range of applications in the field of clinical dentistry, dental research and associated disciplines. One example, cleft lip and palate outcome measurement using MHB is used here to illustrate the use of the technology.

The utilisation of numerical methods for computational geometry and image segmentation on digital dental models and associated algorithm software were key to the development of the first fully automatic landmarking recognition and outcome assessment scoring tool. The machine learning ALR was found to be reproducible and the most accurate and objective method of carrying out dental analyses in the future.

ALR can be applied to both deciduous and permanent dentitions, has widespread applications throughout dentistry, dental anthropology, forensic odontology and in research can constitute an invaluable tool for handling large datasets, eliminating human error and time intensive employment of experts to place landmarks manually. Its implementation promises to provide significant savings in terms of cost and time and an improved patient experience when compared to traditional diagnostic methods.

